# Extracting Biological Insights from the Project Achilles Genome-Scale CRISPR Screens in Cancer Cell Lines

**DOI:** 10.1101/720243

**Authors:** Joshua M. Dempster, Jordan Rossen, Mariya Kazachkova, Joshua Pan, Guillaume Kugener, David E. Root, Aviad Tsherniak

**Affiliations:** Broad Institute of MIT and Harvard, 415 Main Street, Cambridge, MA 02142, USA

## Abstract

One of the main goals of the Cancer Dependency Map project is to systematically identify cancer vulnerabilities across cancer types to accelerate therapeutic discovery. Project Achilles serves this goal through the *in vitro* study of genetic dependencies in cancer cell lines using CRISPR/Cas9 (and, previously, RNAi) loss-of-function screens. The project is committed to the public release of its experimental results quarterly on the DepMap Portal (https://depmap.org), on a pre-publication basis. As the experiment has evolved, data processing procedures have changed. Here we present the current and projected Achilles processing pipeline, including recent improvements and the analyses that led us to adopt them, spanning data releases from early 2018 to the first quarter of 2020. Notable changes include quality control metrics, calculation of probabilities of dependency, and correction for screen quality and other biases. Developing and improving methods for extracting biologically-meaningful scores from Achilles experiments is an ongoing process, and we will continue to evaluate and revise data processing procedures to produce the best results.

## Introduction

Most contemporary cancer drug therapies are broadly toxic to cells. Precision medicine aims to avoid broad toxicity by exploiting cancer-specific vulnerabilities. This approach has shown success with small molecules targeting oncogenes such as B-RAF, ALK, and ER. However, the majority of patients still lack clinically indicated targeted therapies^1,2^. The Cancer Dependency Map aims to address this need and accelerate the development of targeted therapies by charting the landscape of cancer vulnerability^3^. Within this effort, Project Achilles identifies the genetic dependencies of cancer cell lines with over a thousand genome-wide screens, including RNAi knockdown^4,5^ perturbations.

Commitment to open science is a founding principle of the Dependency Map: new data from the Achilles experiment is released to the public every quarter to help drive cancer target discovery. Our procedures for handling Achilles data have evolved as the project continues. However, there is no regular publication associated with Achilles data releases to explain our data processing choices to the public. This document describes in detail the procedures used to turn sgRNA data in Achilles CRISPR-Cas9 screens into the datasets we release. In addition to serving as a comprehensive reference for uses of Achilles CRISPR data, we hope this document contains useful lessons for other groups as they make their own choices about processing CRISPR experiments.

Some new procedures have been used in datasets already public as of the publication of this document; others will be incorporated into future datasets. All the changes described here are scheduled to be in effect by the release of DepMap_Public_20Q1.

## Overview

The Achilles pipeline consists of four stages, named for their outputs: readcount, log fold change, CERES gene effect, and gene dependency (**Fig. 1**). In the readcount stage, deconvoluted sgRNA accounts are assembled and replicates failing fingerprinting, sgRNAs with inadequate pDNA representation, sgRNAs with suspected off-target activity, and replicates with too few reads are removed. At the log fold change stage, individual replicates and whole cell lines with inadequate control separation or poor replicate correlation are removed, then CERES inference is performed. As part of CERES, sgRNAs are aligned to the hg38 genome assembly and sgRNAs with no annotated gene targets are dropped. In the CERES gene effect phase, the CERES output is rescaled and corrected to remove known confounders and common dependencies identified. Finally, for each gene score in each cell line we infer the probability that the gene’s score represents a true biological depletion phenotype.

**Fig. 1:**
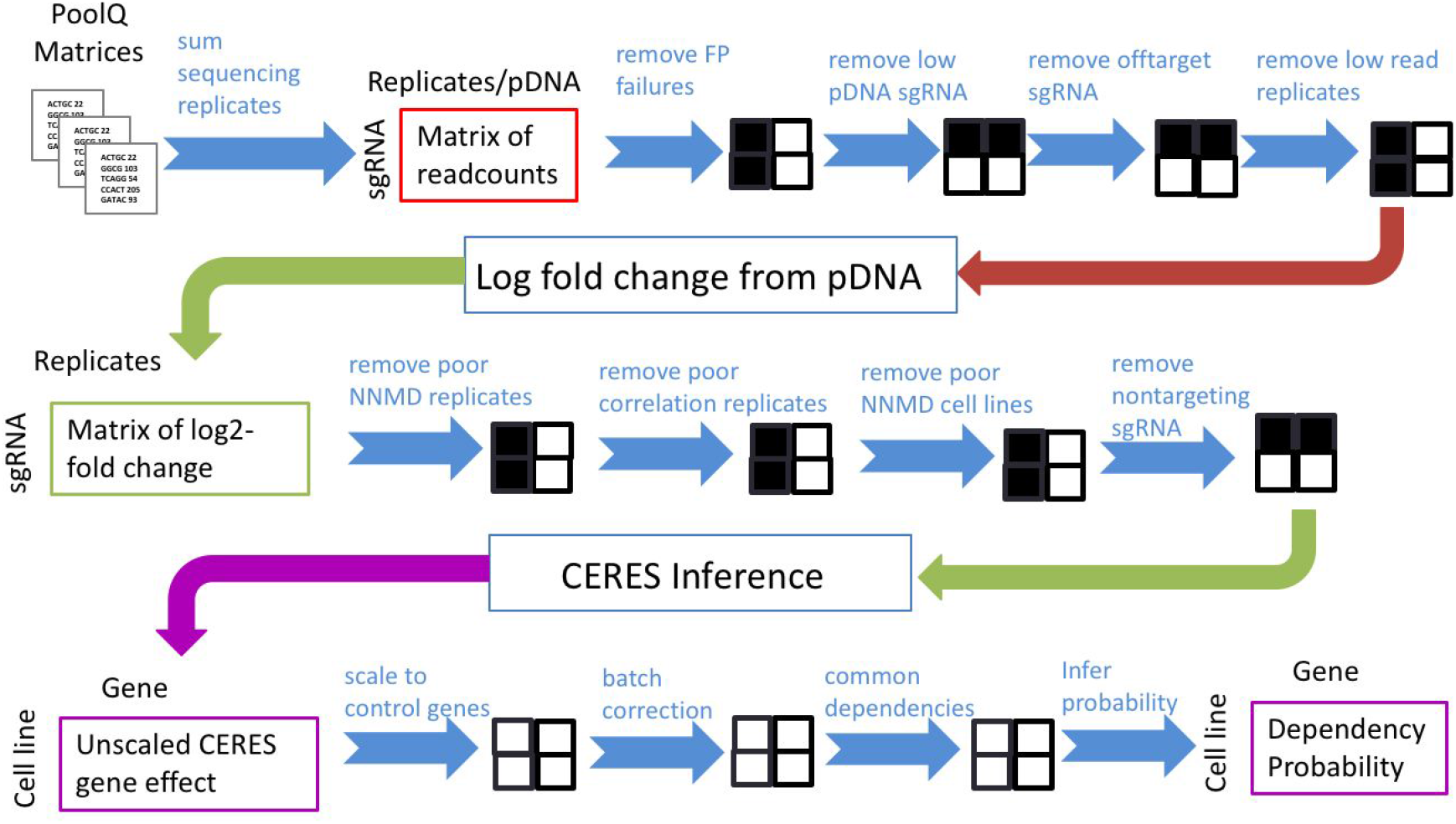
The Achilles data pipeline. The major steps used to generate Achilles matrices from deconvoluted screen sgRNA readcount data. This pipeline applies to public 19Q1 and later datasets. Black and white grids illustrate which axis is being reduced during QC steps.

Various details of this pipeline have evolved over time. Changes are described throughout this document. For reference, **Table 1** lists the changes introduced with each quarterly release.

**Table 1:** Changes introduced in the Achilles pipeline by release date.

| Release | Changes |
| --- | --- |
| Public 18Q1 | Introduced gene_dependency, pan_dependent_genes, and a standardized file naming system and formatting;<br>Added Entrez IDs to gene names;<br>Identified suspected off-target guides and removed them |
| Public 18Q2 | Changed copy number prioritization from preferring Broad data of all kinds to preferring WES data from all sources;<br>Removed compound gene annotations. |
| Public 18Q4 | Changed cell line indexing to use DepMap IDs (called Broad IDs in this dataset only);<br>Changed QC to drop individual replicates that show insufficient control separation before conducting cell line QC |
| Public 19Q1 | Introduced batch corrected versions of gene_effect, gene_dependency, and pan_dependent_genes |
| Public 19Q2 | Made batch correction the default form of the data;<br>Began providing gene_effect_unscaled (the raw CERES output);<br>Public releases combined with CCLE releases;<br>Achilles file names have the prefix "Achilles_" added |
| Public 19Q3 | Changed reference assembly to hg38;<br>Changed positive controls used for scaling the data to a more inclusive and recent list;<br>Changed gene dependency probability inference to use the same positive controls used for scaling, rather than those found from the data itself;<br>Changed cell line and replicate QC metrics to use NNMD for control separation and pDNA-normalized Pearson correlation of high-variance genes for replicate reproducibility; |
|  | Changed batch correction to only remove five principal components from the data;<br>Retained sgRNAs failing QC in logfold_change and provide mapping information for them in *Achilles_dropped_guides.csv*;<br>Removed dropped_guides<br>Removed Y chromosome data from gene_effect_unscaled and later files |
| Public 19Q4 | None anticipated. |
| Public 20Q1 | Other changes undergoing evaluation |

## Data Generation and Assembly of Read Counts

Replicates from the Avana library are sequenced in four technical replicates and deconvoluted as described in Meyers *et al*.^5^ Read counts of single guide RNAs (sgRNAs) from technical replicates of the same cell line are summed together. Additionally, with each sequencing run the Broad resequences the primary DNA (pDNA) pool of Avana sgRNAs. The Broad has adopted four different processes for PCR amplification over the lifetime of the Achilles CRISPR experiment with transitions occurring in July 2015, November 2015, and April 2016, yielding four pDNA batches numbered 0-3. Batches for each replicate and pDNA measurement are provided in *replicate_map*.

Read count measurements belonging to the same batch are summed and combined. All biological replicate and pDNA batch read counts are combined in a read count matrix (*Achilles_raw_readcounts.csv*) with replicates and pDNA batches as columns and rows corresponding to the total number of sgRNA reads.

We apply four filters to the resulting read count matrix. First, we remove suspected off-target Avana sgRNAs. These sgRNAs were identified from early runs of CERES that identified a single sgRNA from among those targeting a gene as the sole efficacious sgRNA for that gene. A list of these sgRNAs is provided with each Avana dataset as (*Achilles_*)*dropped_guides.txt* for data sets before 19Q3. In later datasets, failed guides also have their gene targets indicated in the file *Achilles_dropped_guides.csv*. The sgRNA abundance in the pDNA pool is not uniform but roughly log-normal as expected of the results of an exponential process, with a tail of sgRNAs with low or zero representation (**Fig. 2**). Steps contributing to the range of pDNA sgRNA abundance include the production of oligos on chip, DNA cloning, and amplification in bacteria. To prevent excessive noise, we set to null (NA) pDNA measurements for sgRNAs if the measurement is less than one read per million for that pDNA batch, corresponding to about 0.605% of all sgRNA reads out of 74,661 sgRNAs (**Fig. 2**). sgRNA reads are then NAed in all replicates belonging to the same pDNA batch. Next, we remove all replicates that fail to match their parent cell line using SNP fingerprinting (described further below). Finally, we remove replicates that have fewer than 15 million total read counts. These are sent for resequencing to generate sufficient read depth.

**Fig. 2:**
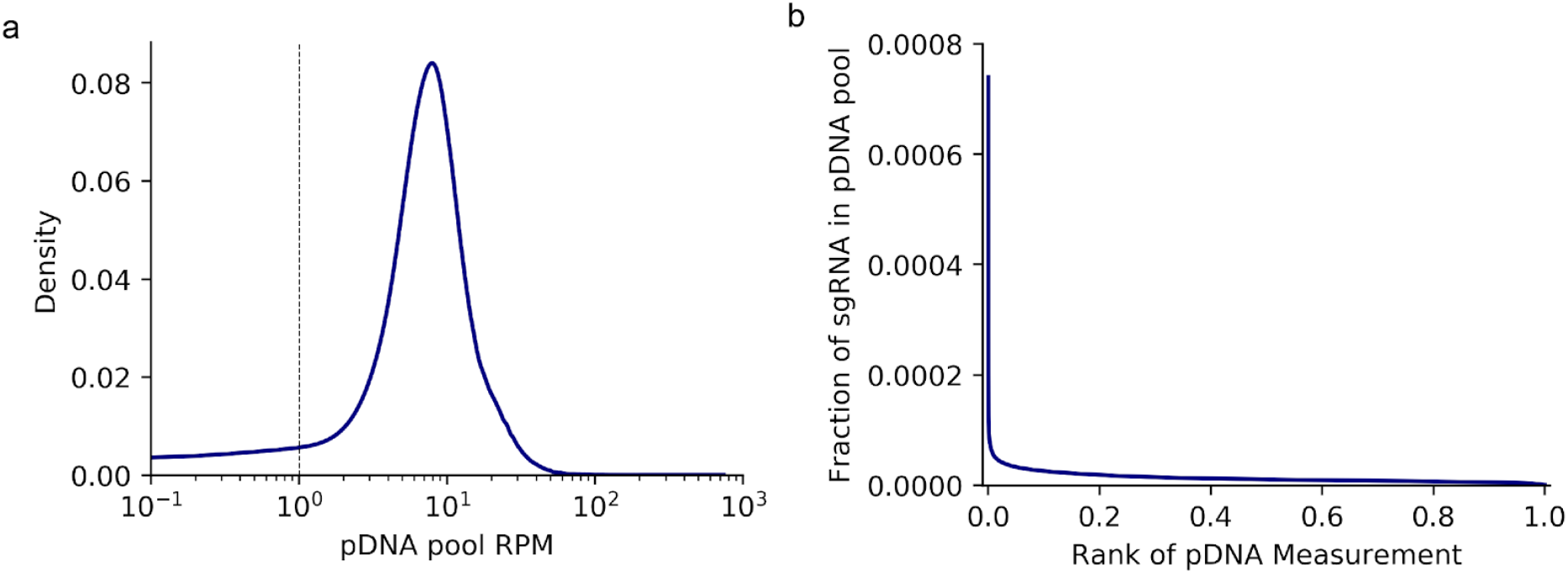
pDNA distribution. (a) The distribution of pDNA reads per million. The dotted line indicates the threshold below which sgRNA reads will be NAed. (b) Fraction of mapped pDNA pool reads that map to individual sgRNAs in descending order.

We normalize each read count column by its total reads, multiply by one million, and add one pseudocount to generate reads per million (RPM). We take the log2 of the ratio of each replicate RPM to the corresponding pDNA batch RPM to generate an sgRNA by replicate logfold change (LFC) matrix.

## Data Quality Control

Quality control for the Achilles CRISPR data proceeds in the following order: (1) the fingerprint integrity of each replicate is checked and failing replicates are dropped, (2) replicates are next dropped if they show insufficient separation between positive and negative control genes, (3) surviving replicates are dropped if they are not sufficiently similar to other replicates of that cell line, and (4) LFC values of the remaining replicates of a cell line are averaged and control separation is once again checked. Cell lines failing the QC process are not included in the released dataset.

### Fingerprinting

Prior to data being put through the Achilles pipeline, it is necessary to ensure that no contamination, sample swapping, drastic mutation, or other integrity-compromising events occurred during the screening process. We control for these events by checking that the “fingerprint” of each sample (96 SNP calls uniquely identifying a cell line) matches to the original cell line both after cas9 infection and after the gene knockout is performed. We calculate the Pearson correlation between samples by transforming each SNP call into a numerical value using a SNP reference file with the following logic: assign 1.0, 0.5, 0.0 to each SNP call if it is homozygous for the alternate allele, heterozygous, homozygous for the reference allele, respectively^6^. In the case of aneuploidy, a homozygous SNP would not affect the SNP calling, but a heterozygous SNP would cause a failed call due to unequal signal from alleles; a sample with insufficient called SNPs will fail fingerprinting. For samples A and B to match, the Pearson correlation between the SNP calls of A and B must be greater than or equal to 0.9, and the z-score of the Pearson correlation between A and B must be greater than or equal to 3.0 relative to the distribution of Pearson correlations between A and every sample that we have seen so far. With this metric, we check that the fingerprints of the Cas9-positive “derivative” samples and sgRNA-infected “screen” samples match the fingerprint of the original “parental” cell line. 95% of screens match their respective parental (***Fig. 3***). Additionally, the p value for a screen matching a parental other than its own given the distribution of z-scores of unpaired parentals and screens is 0.00038 (***Fig. 3b***). In addition to excluding contaminated samples, this method allows us to identify and reverse labeling errors. In one case, we found that the derivative for DMS53 matched the parental for HCC38 rather than its own (and vice versa), indicating that either parental or derivative data had been swapped.

**Fig 3:**
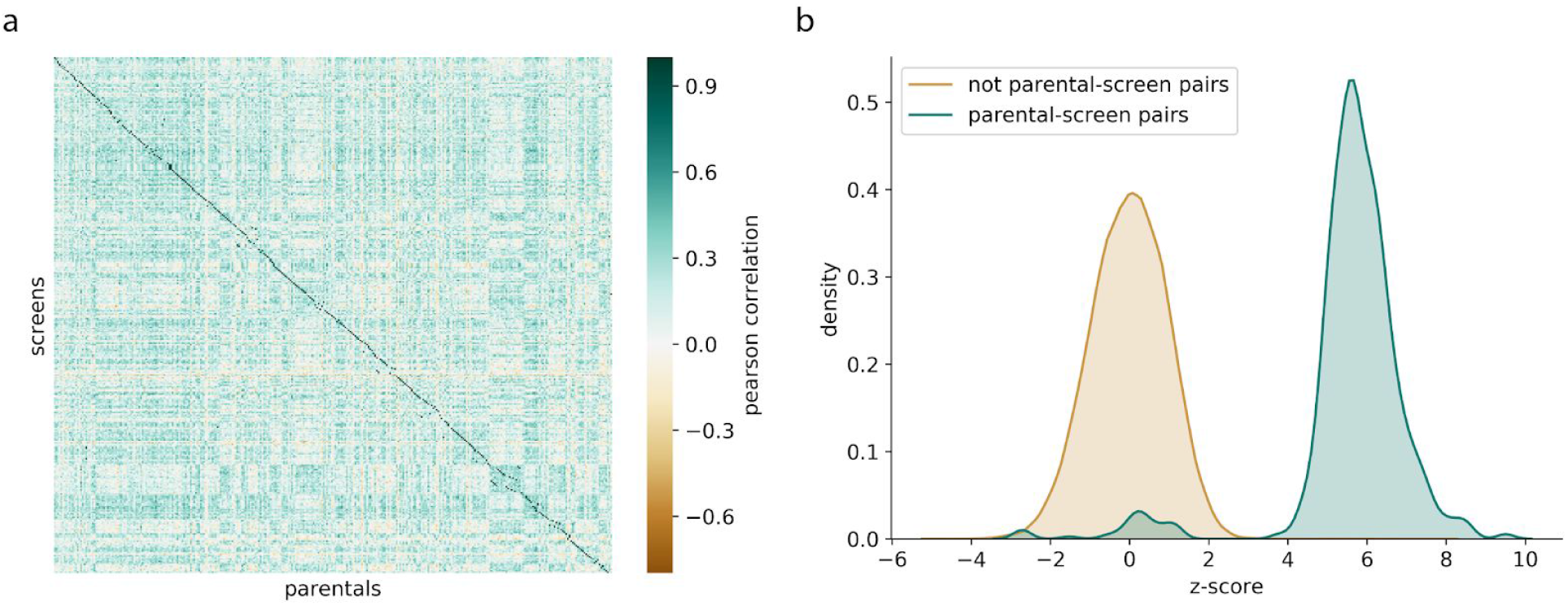
Visualization of metrics for fingerprinting: (a) Correlations between SNP calls of screens and parentals with matched parental-screen samples on the diagonal. (b) Distribution of z-score values of parental-screen pairs and non parental-screen pairs.

### Choice of Gene Controls

For quality control and normalization, we use exogenously defined nonessential genes as negative controls and common essential genes as positive controls. The nonessential gene list was taken from the study by Hart *et al*. in RNAi data^7^ (*nonessentials.txt*). The essential gene list is the intersection of two studies^8,9^ that use orthogonal methods, gene trap and CRISPR-ko, to make a discrete list of genes required for the life of a human cell line. Blomen *et al*. suggest using the overlap of hits in KBM7 and HAP1^8^. Hart *et al*. suggest the “daisy” method of using hits that overlap at more than half the samples, where a hit is defined as having a BF > 0 ^9^. Taking the intersection of the two studies results in 1,248 cell line essential genes reported in *common_essentials. txt*.

### Separation of Gene Controls

The separation control step in the Achilles pipeline is used to check that essential and nonessential genes are behaving as expected. Previously, we used the strictly standardized median difference (SSMD) to assess the separation between the killing effects of the two groups. However, SSMD penalizes high variance in the LFC values of both the essential and nonessential genes. While we expect the LFC values of the nonessential genes to have a tight distribution centered around zero, no analogous expectation exists for the essential genes. Instead, we measured the difference between the means of the LFC values of the essential and nonessential genes divided by the standard deviation of the LFC values of the nonessential genes, which we refer to as the null-normalized mean difference (NNMD; ***Fig. 4a***). Note that for both NNMD and SSMD more negative scores indicate a better screen.

**Fig. 4:**
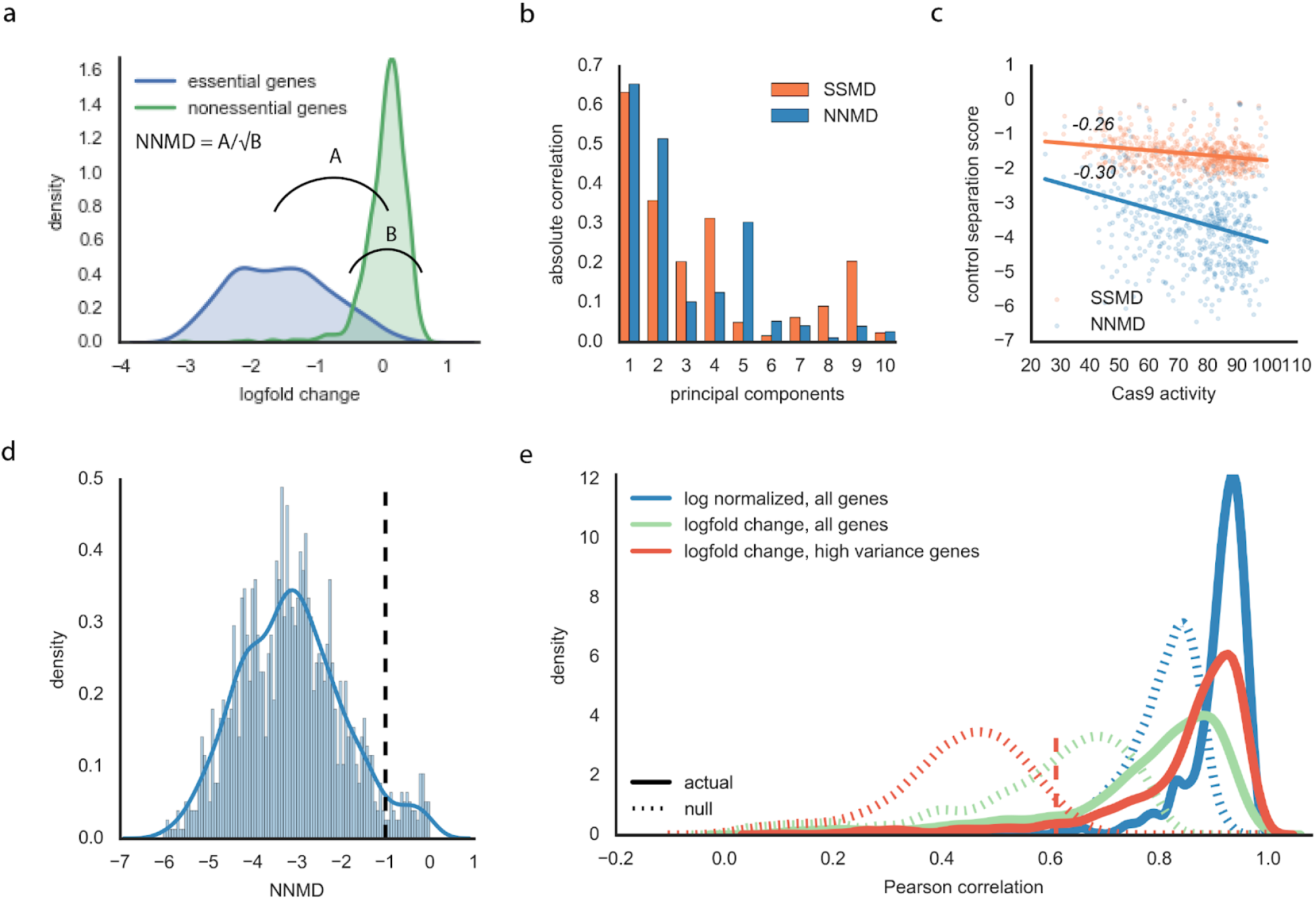
Changes to quality control metrics: (a) Visualization of NNMD for a single replicate; A represents the difference in means between the essential and nonessential genes, B represents the variance of the nonessential genes. (b) Absolute Pearson correlation between each of the first 10 principal components of the replicate LFC values and SSMD and NNMD. (c) Scatter plot showing SSMD and NNMD relationship to Cas9 activity. (d) Distribution of NNMD scores of all the replicates. Line at −1 indicates the new separation control threshold. (e) Null and actual distributions for various data. Dashed red line represents 0.05 p-value cutoff using the null distribution for LFC values, high variance genes; this is the replicate reproducibility threshold.

NNMD is more strongly related to the first two principal components of the data (absolute Pearson’s r = 0.65 and 0.52, respectively; ***Fig. 4b***) as well as Cas9 activity (Pearson’s r = −0.30; ***Fig. 4c***) than SSMD (absolute Pearson’s r = 0.63,0.36; Pearson’s r = −0.26, respectively). Both Cas9 activity^5^ and the first principal components (see Batch Correction) are strongly related to screen quality, suggesting that NNMD is a better quality metric. We require a threshold of −1.0 for NNMD based on a local minimum in the distribution of the NNMD scores across all replicates (***Fig. 4d***).

### Agreement between Replicates

Because at least two replicates were performed for each screen, we computed the Pearson correlation between the replicates to ensure that the results are reproducible. Previously, we computed the Pearson correlation using log2-normalized read counts using all genes.

However, we observed that using log2-normalized data can produce high Pearson correlations between random pairs of replicates driven by the pDNA distribution alone; we instead used gene-level LFC values instead as these are more sensitive to cell-line specific screen response. Additionally, when calculating correlation, we restricted genes to those with high variance across cell lines identified from the top 3% most variable scores in the public 18Q4 gene effect (*Achilles_high_variance_genes.csv*). The separation between the null distribution and actual distribution of Pearson correlations between replicates is larger using high variance genes than all genes, meaning we better capture a distinct representation of each cell line using high variance genes (***Fig. 4f***). We require a Pearson correlation threshold of 0.61, equivalent to a *p-value* of 0.05 for observing the correlation between random pairs of replicates. In the case of more than two replicates per cell line, replicates not reaching the 0.61 threshold when correlated with any of the other replicates for that cell line are excluded from the dataset.

## Estimation of gene effect scores

Cas9 cutting of DNA induces a toxic effect that increases with the number of sites cut. This results in the “copy effect,” in which sgRNAs targeting highly amplified regions of the genome produce depletion effects comparable to knockout of an essential gene even when the specific locus targeted is an intergenic region^10^. To remove the copy effect, the Broad Institute developed the CERES package^5,10^. Using CERES consists of two phases: prepare_inputs, which aligns sgRNAs to genes and identifies the number of cuts expected for each sgRNA in each cell line; and run_ceres, which treats each observed sgRNA log fold change as the sum of a gene and copy number effect, multiplied by a guide efficacy. More details can be found in Meyers et. at.^5^.

### Alignment of sgRNAs to genes

In previous Achilles releases, sgRNAs were aligned to hg19. Beginning with DepMap_Public_19Q3, sgRNAs will be aligned to hg38. Alignments are performed using Bowtie to find all exact matches to the 20-nt sgRNA sequence. Matches are then filtered for the presence of a NGG PAM sequence at the 3’ end.

Matches which fall in the exon of a gene as annotated by the Consensus Coding Sequence (CCDS) GRCh38.p12 (GRCh37.13 for Achilles releases DepMap_public_19Q2 and earlier) are annotated as targeting that gene. CERES models multiple gene targeting as linear and additive. This assumption can fail for synergistic cases, as when a single sgRNA targets many members of a family^11^. It can also introduce problems in the case of compound genes (those containing the superset of two other genes’ transcripts). With compound genes, CERES can arbitrarily shift the scores of the two individual genes in one direction (negative or positive) while shifting the scores of the compound gene in the opposite direction. The resulting scores produce good matches to the sgRNA log fold change data according to the CERES model, but the inferred gene effects are unlikely to be biological. As it is uncertain how often the compound forms of most genes really occur in cells, we have opted to avoid this problem by removing compound genes from gene annotations from avana_public_18Q2 onwards.

### sgRNA Copy Number

In previous releases of Achilles, it was also necessary to discard genes located on the sex chromosomes. This was due to systematically biased copy number estimations for these chromosomes using the single-nucleotide polymorphism (SNP) pipeline, which normalized copy number to the background across cell lines. Combined with CERES this led to small but persistent biases in the scores for genes located on these chromosomes. This problem does not arise with WES-based copy number. Additionally, there is clear evidence that WES copy number has a generally stronger relationship with log fold change than SNP copy number. The Broad has therefore undertaken whole-exome sequencing of all cell lines in Achilles that have not previously been sequenced by Sanger. Beginning with avana_public_19Q3, cell lines with WES-based copy number were released with CERES scores for the X chromosome, and by 20Q1 we anticipate that all cell lines will have WES copy number and X chromosome scores.

In the interim, in cases where cell lines have copy number (CN) calls from multiple sources, we use the following priority rules: (1) Broad WES-based CN calls are always used if available. (2) If a cell line has both Sanger WES-based and Broad SNP-based calls, we run a correlation analysis using Achilles LFC to determine which should be used. This analysis addresses the possibility of differences in the CN profile due to cell line heterogeneity that arises as a result of culturing, as described by Ben-David *et al*.^12^. As several cell lines have been shown to have distinct CN profiles between the Sanger and Broad Institutes due to these effects, we run this analysis to determine if the Sanger WES-based CN profile can be used for each cell line.

In general, the correlation analysis we introduced above uses the gene-level relative CN profile. Relative CN for each profile is multiplied by 2 to be relative to a ploidy of 2 and capped at a maximum value of 20 for each gene in the Achilles screen. The Pearson correlation is calculated between the Achilles LFC and CN profile from the two sources. Additionally, we calculate the number of relative CN calls where the difference in log2(relative CN) between the two sources is greater than 0.5. This gives us a measure of dissimilarity between the two samples, where a higher percent difference indicates more dissimilarity between the calls. When prioritizing the CN profile to use for a cell line with Sanger WES and Broad Institute SNP profiles but no Broad Institute WES profile, the Sanger WES profile is used unless this profile has a more positive Pearson correlation value and has a percent difference greater than 2.5% with the Broad Institute SNP profile. For example, MFC7 is a cell line for which it was determined that the Broad Institute and Sanger were working with different clones. Examining MCF7’s two CN profiles, we observe distinct CNAs between the two centers (**Fig. 5a**). When we correlate its gene-level LFC with the CN profiles, we observe a more negative correlation with the Broad Institute SNP profile over the Sanger WES profile (**Fig. 5b**). Based on this analysis, the Sanger WES profile is too dissimilar to the cell line screened in Achilles and should not be used for CN correction. 25 cell lines failed this correlation analysis from DepMap_public_19Q1 (**Fig. 5c**).

**Fig. 5:**
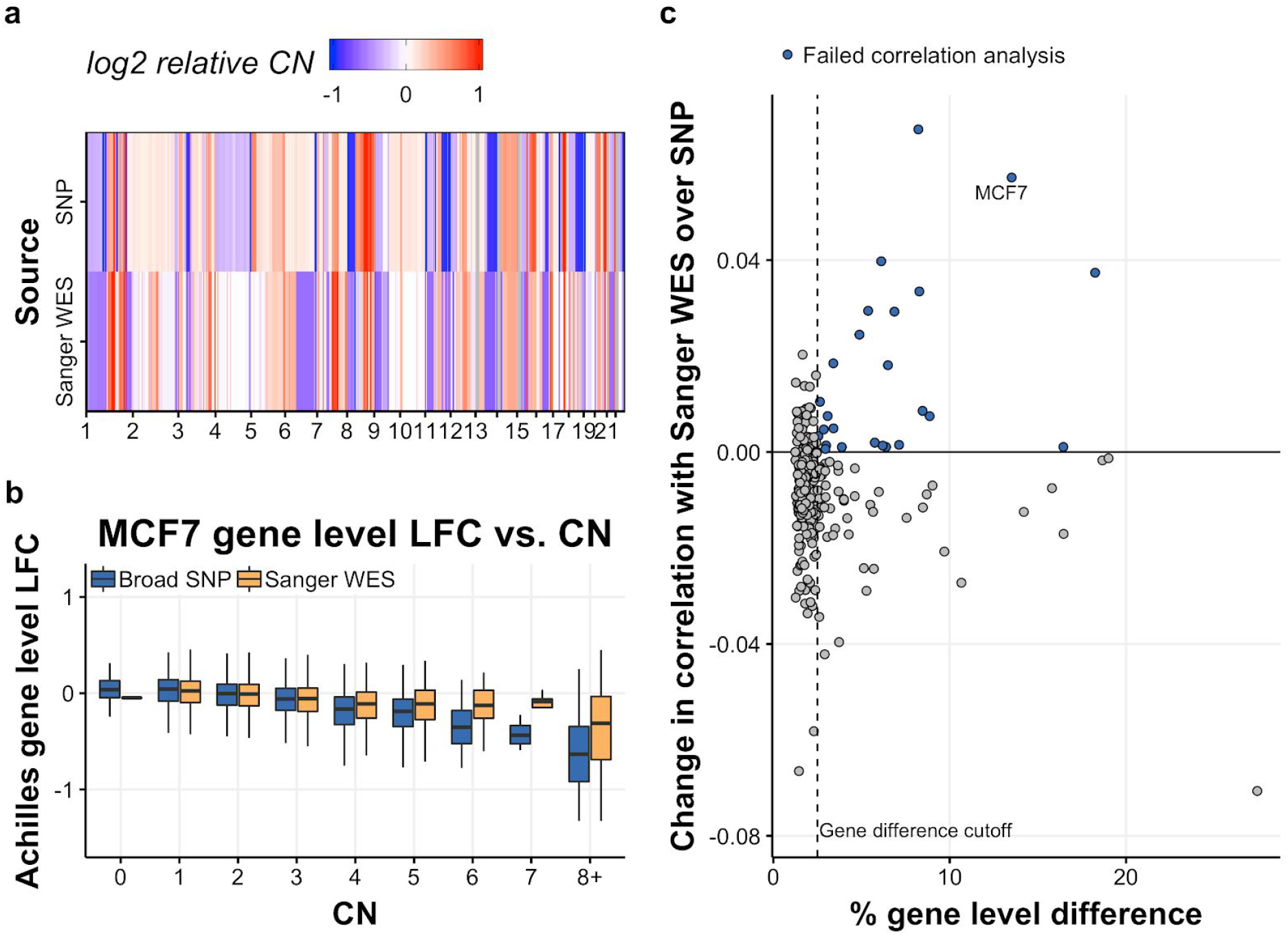
Copy number profile prioritization: (a) Sanger WES and SNP-based CN calls across the genome for MCF7. (b) Achilles gene-level LFC vs. relative CN for MCF7. CN calls are binned into integer bins and the distribution of LFC from the Achilles screen is plotted. (c) Change in correlation when using Sanger WES-based CN calls over SNP-based CN calls vs. the % of genes with a log2(relative CN) difference > 0.5. The vertical dashed line indicates the passing cutoff used for % gene-level difference. The horizontal black line at 0 indicates where a cell line would fall if it had the same Pearson correlation when using Sanger WES and SNP-based calls. Cell lines that failed the correlation analysis are blue.

### Running CERES

The CERES method is described in Meyers et. al.^5^, and the code is available at Github^13^. The critical hyperparameter of the CERES model is lambda_g, which controls the strength of the regularization of individual gene scores towards the mean. A high value for this parameter produces good separation of common essential and nonessential genes, at the expense of compressing the scores of selective essential genes. We have chosen the value 0.4 as the compromise between these competing considerations. This value is slightly less than the values that tend to maximize out-of-sample accuracy in most datasets. However, because most genes are either essential or nonessential in all lines, maximizing global out-of-sample accuracy will produce over-regularized datasets for the purpose of identifying selective dependencies. CERES-estimated gene effect scores are provided unaltered (except for filtering sex chromosome scores in lines with SNP copy number estimates) as *Achilles_gene_effect_unscaled.csv*. Estimates of sgRNA efficacy and offset are given in *Achilles_guide_efficacy. csv*.

### Normalization

CERES scores are an abstract quantity without direct biological interpretation. Additionally, common essential genes exhibit high mutual correlation driven by the technical quality of the Cas9 screen in cell lines. To improve interpretability and reduce technical confounding, we shift and scale the scores in each cell line so the median of negative controls is 0 and the median of positive controls is 1. Previously we used essential and nonessential genes identified in RNAi screens for this purpose. However, we find that the positive control genes identified this way tend to be among the most extreme responders. Beginning with DepMap_public_19Q3, Achilles releases will use a list of common essential genes found from CRISPR screens. This will create a change in the scale of gene_effect in 19Q3, amplifying all values by about 30%.

## Correction of Experimental Confounders

### Bias Related to Experimental Confounders

A common problem in high-throughput experimental data is the separation of biological signal from systematic biases. In the Avana dataset, changes over time in the technology used to perform the screen and differences in the compatibility of cell lines with the assay create variation that is unrelated to gene dependency. These factors, specifically the pDNA batch and Cas9 activity of a cell line, along with two measures of screen quality, NNMD and replicate correlation, are referred to as experimental confounders. Together, experimental confounders explain 11.9% of the variance in scaled, uncorrected gene effect data.

To further assess the extent of data contamination by experimental confounders, we perform principal component analysis (PCA) on gene effect level data and study the relationship of the dominant components to confounding features. We are able to create strong predictive models for the first six principal components using only experimental confounders. Combined, these principal components explain more than 20% of the dataset-wide variance. A similar bias is present at all levels of processing, suggesting the phenomenon is a result of experimental design, not a specific data processing step (**Fig. 6a, 6b**).

**Fig. 6:**
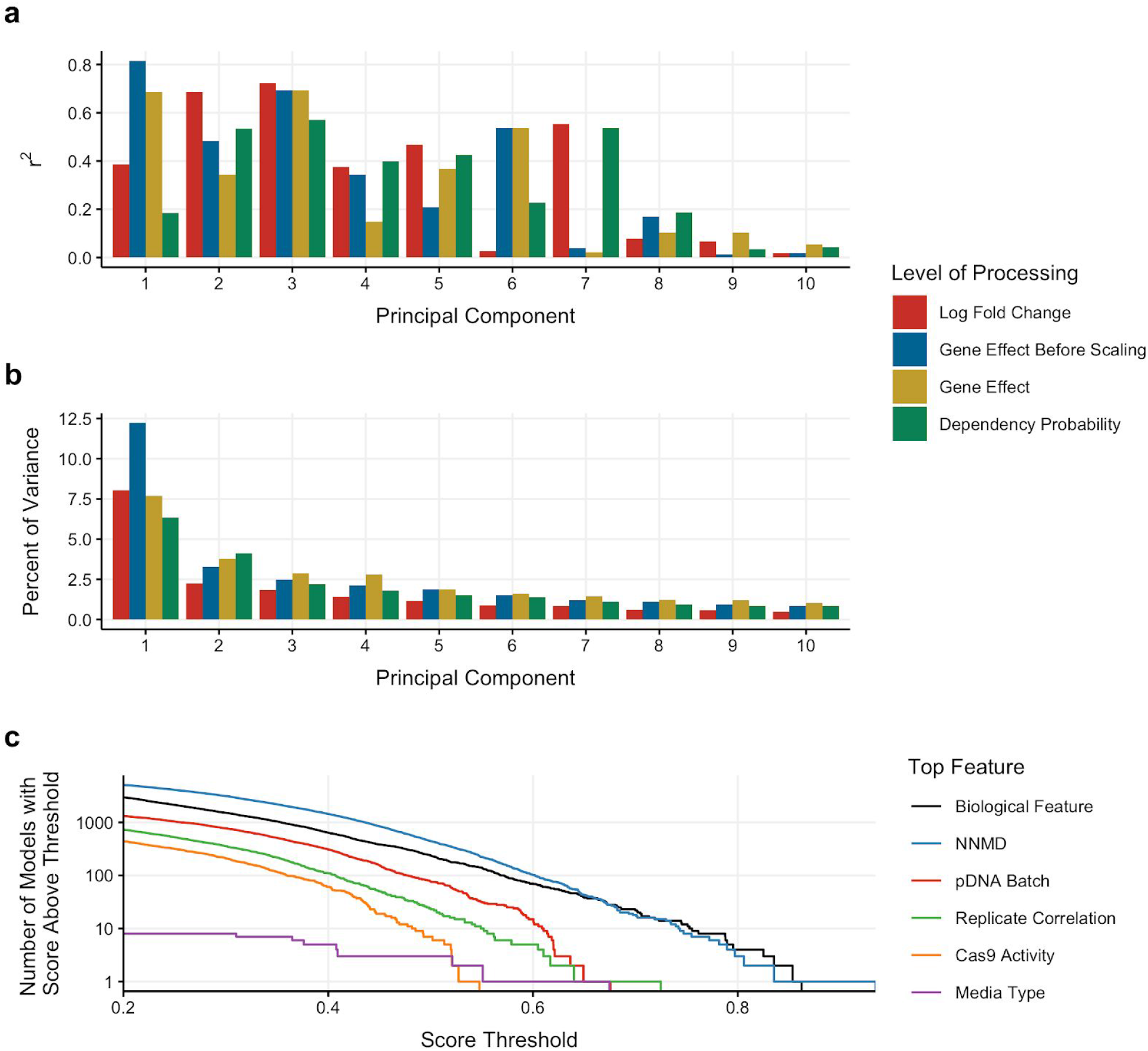
Impact of technical bias on Avana data: (a) Performance of a linear model trained using pDNA batch, cas9 activity and NNMD to predict PCs 1-10 after each primary level of processing. (b) Percent of variance explained by PCs 1-10 after each primary level of processing. (c) Survival curves for models trained to predict gene knockout effects using CCLE cell line features and experimental confounders. Only models with score > .2 are shown. The “score” of a model is defined as the Pearson correlation between a gene’s observed and predicted dependency profiles out of sample.

Additionally, we find that experimental confounders are often stronger predictors of individual gene scores than biological features. We construct predictive models for the gene dependency scores of each gene using a combination of experimental confounders and CCLE cell line characterizations as features. (Analysis Methods). To evaluate model performance, we define the “score” of a model to be the Pearson correlation between a gene’s observed and predicted dependency profiles out of sample. More than 61% of trained models identify an experimental confounder as their top feature. This result is especially striking as experimental confounders make up only 17 of 165,209 features supplied to the model. Furthermore, given any score threshold less than 0.67, the majority of models with scores exceeding the threshold use a technical artifact as their top feature (**Fig. 6c**).

In the following three sections, we will first describe a new step in the Avana data processing pipeline called Removal of Confounding Principal Components (RCPC). Next, we will compare RCPC to alternative methods on the basis of their ability to remove unwanted variation and denoise biological signal. In the last section, we will examine the effects of RCPC on dependency prediction.

### Description of Method for Correction of Experimental Confounders

Many solutions have been proposed for separating unwanted variation from biological signal. We choose a PCA-based denoising procedure because of its simplicity and effectiveness. PCA captures systematic variation that exists across the dataset, while discounting gene-specific information. RCPC is implemented as follows: PCA is performed on gene-effect level data and the first six principal components and component loadings are removed. The truncated principal component and component loading matrices are multiplied to obtain a matrix of the original dimensions. Cell lines are rescaled so that the median negative control gene effect score for each cell line is 0 and the median positive control gene effect score for each cell line is −1. This procedure is applied to public releases starting with avana_public_19q1, where both corrected and uncorrected versions of the data are provided. In later releases, post-CERES data is corrected by default. The batch-corrected CERES gene effect scores are given in *Achilles_gene_effect.csv*.

### Comparison with Alternate Methods

We limit the scope of our comparison to methods which do not require specification of a per sample phenotype and which preserve the absolute scale of the gene effect level data (where a single score indicates the dependency level of a cell line, not the dependency of a cell line relative to all other cell lines). Methods which require specification of a phenotype were not considered for two reasons. First, corrections that utilize phenotype have the potential to introduce misleading results^14^. Second, the large number of use cases for the Avana dataset make the choice of specific phenotypes difficult. Given these constraints, we present comparisons of RCPC to several methods: a linear-regression based approach, two previously published methods for isolation of confounders in the context of RNAseq data (ComBat, RUV2)^15,16^.

In addition to the above, we also investigate a recently developed method^17^ which utilizes a novel correction of the Avana dataset to identify co-functional gene interactions. The authors use PCA on a set of negative control genes to isolate undesirable systematic variation. In this report, we refer to this method as negative-control PCA correction (ncPCA). The original ncPCA method is not appropriate for the standard Avana data release because it does not preserve the absolute dependency status of genes: after correction, gene means approach zero. However, we developed an adaptation of ncPCA which we call modified negative control PCA correction (ncPCA mod). For details on how the methods are applied, see the Analysis Methods section.

All presented methods remove the observed relationship with experimental confounders (**Fig. 7a**), so correction methods are also evaluated according to their ability to improve biological signal according to four criteria: (1) strength of associations between known pairs of related genes (**Fig 7b**); (2) agreement of cell line clustering with tissue of origin annotations (**Fig 7c**); (3) strength of association between gene dependency data and gene expression data (**Fig 7d**); (4) strength of known biomarkers of dependency (**Fig. 7e**). All of the presented methods yield substantial improvements in metrics 1-3, but cause decreases in the strength of biomarker-gene associations for a set of 25 genes with strong, consistent, biomarkers across the DepMap CRISPR knockout and RNAi datasets. These biomarker-gene associations are referred to as “consensus biomarkers” (**Supplementary Table 2**). RCPC outperforms the other methods by most metrics, but was not the top choice with respect to two metrics: RCPC yields a smaller increase in the number of discovered relationships between known paralog genes compared to two other methods. RCPC also caused the second-largest decrease in the strength of Spearman correlation for consensus biomarkers; RCPC decreases the median Spearman correlation from 0.55 to 0.50.

**Fig. 7:**
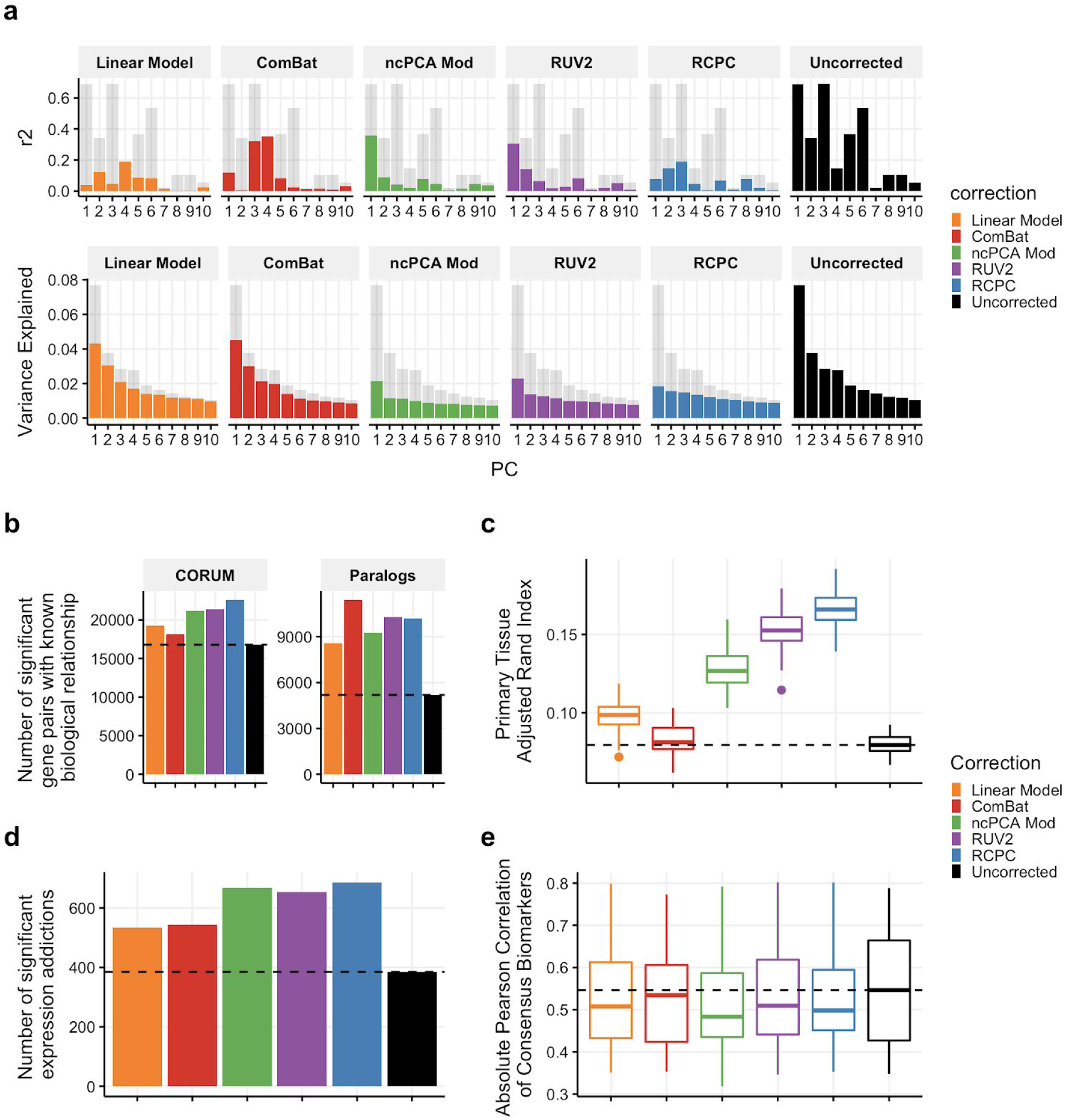
Comparison between correction methods: (a) Top: Performance of a linear model trained to predict linear components of corrected gene effect data using experimental confounders. and the percent of variance explained by each principal component. Bottom: The percent of variance explained by each principal component. (b) The number of related gene pairs with a statistically significant Pearson correlation. Left: Gene pairs are components of the same protein complex (annotated by CORUM). Right: Gene pairs are paralogs (annotated by Ensembl). (c) Agreement between cell line primary tissue type-based group assignments and cell-line clustering by sequential t-SNE and k-means. (d) The number of genes with statistically significant negative correlations between the gene’s Avana knockout profile and CCLE gene expression profile. (e) Dependency-biomarker Pearson correlations for consensus biomarkers. Consensus biomarkers are genes with strong predictive models that are consistent across CRISPR KO and RNAi datasets.

The ncPCA method has been reported to show large gains in identifying functional gene relationships^17^. Accordingly, we compare ncPCA and RCPC based on their utility for discovering co-functional gene interactions and their ability to remove systematic bias. We observe that while ncPCA reduces systematic bias among nonessential genes, it reimposes this bias on essential genes. This phenomenon is exemplified by PC1, which is identified as explaining the majority of the bias among the negative control gene set^17^. After ncPCA, correlations to the loading of PC1 decrease among nonessential genes but increase among essential genes (**Fig. 8a**). The introduction of a confounding effect results in inflated correlations among all essential genes regardless of biological function (**Fig. 8b**). In the original publication of ncPCA, a 50-fold increase in the number of significantly correlated genes is reported. This is partially a result of the introduction of a shared confounding effect and partially a result of an improvement in biological signal quality. We use a previously developed method^18^ for evaluating the significance of protein complexes in fitness similarity networks to compare the utility of RCPC and ncPCA for discovery of gene-interactions and find that RCPC outperforms ncPCA in this regard (**Fig. 8c**).

**Fig 8:**
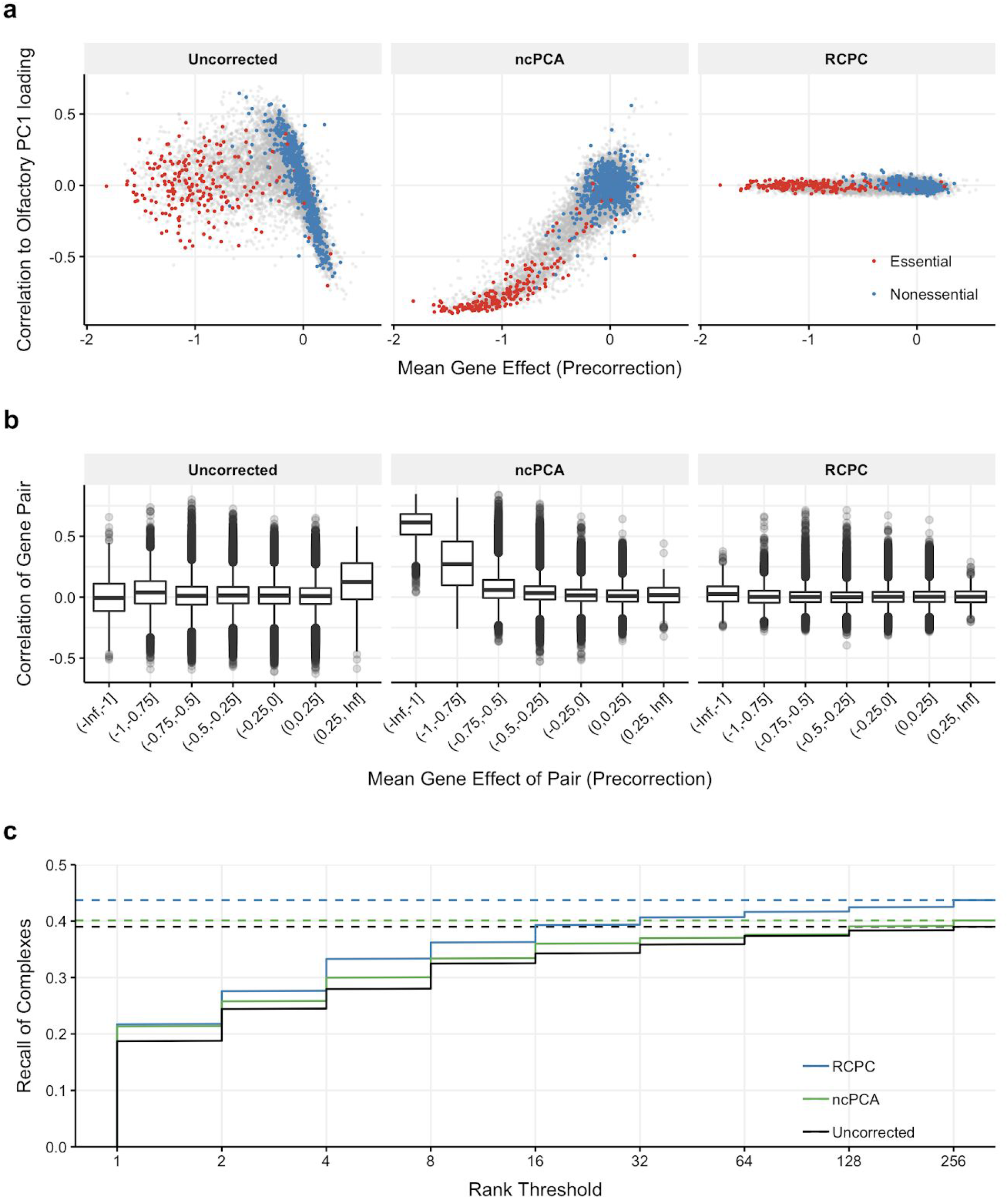
Effect of ncPCA: (a) Effect of correction on PC1 related bias. In the uncorrected data, nonessential genes exhibit a strong association with PC1 of the olfactory genes. After application of the ncPCA, PC1 related bias has been transferred from nonessential genes to essential genes. (b) Effect of correction on gene pair correlations. Three million gene pairs were randomly selected for visualization. (c) Recall of CORUM complexes with a significant internal-to-external edge ratio colored by correction method.

To illustrate the confounding effect of ncPCA, we draw networks containing four essential complexes and include the top 400 ranked correlations (within or between complex subunits) as edges. For the uncorrected and RCPC datasets, the majority of edges lie within protein complexes. In contrast, after ncPCA, most edges are between essential genes with high mean effect score in different complexes (**Fig. 9**). Because ncPCA creates indiscriminately high correlations between most essential genes, we recommend the use of RCPC.

**Fig. 9:**
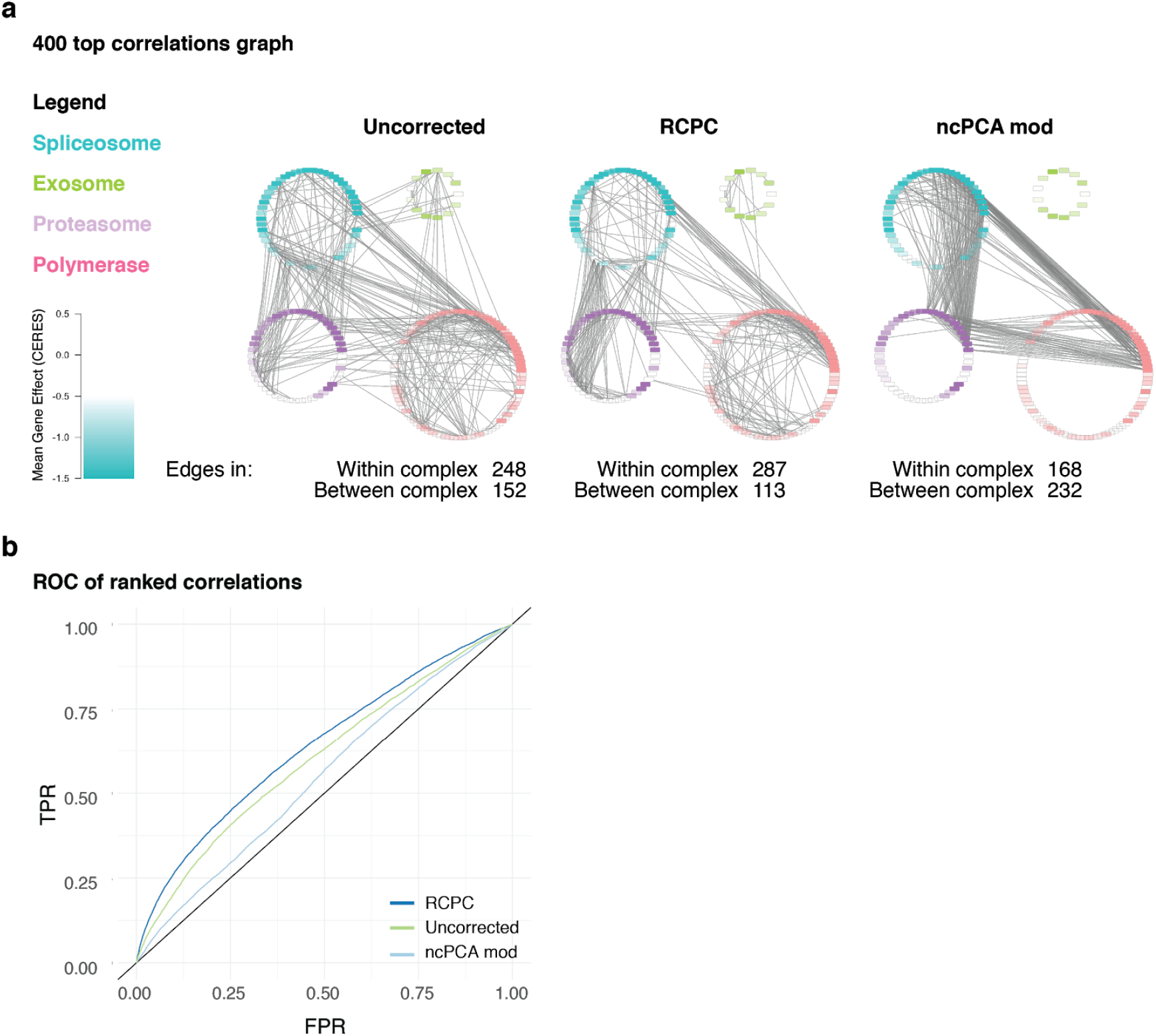
Visualization of the confounding effect from ncPCA: (a) Network visualization of the top 400 strongest correlations within the splicesome, exosome, proteasome and polymerase complexes. Nodes are colored by mean gene essentiality. (b) ROC curve for the spliceosome, exosome, proteasome and polymerase complexes when all edges above a given correlation rank for each gene are considered, only considering correlations that exist between genes in one of the depicted complexes.

### Effect of Correction on Dependency Predictions

One of the primary use cases of the Avana dataset is dependency prediction. To identify biomarkers of dependency, we construct predictive models for the gene effect scores of each gene using a combination of experimental confounders and CCLE cell line characterizations as features (Analysis Methods). To quantify the effect of the correction method on dependency prediction, we compare the performance of models trained on uncorrected data to that of models trained on RCPC corrected data. The impact of confounders is almost completely eliminated (**Fig. 10a**). After correction, the percentage of models that use a technical artifact as their top feature decreases from 61% to 1%. This improvement holds at all levels of performance. At most, 3% of models with scores greater than any threshold use a technical artifact as their top feature. While systematic bias is reduced, we also observe an overall decrease in model predictability. Model predictability decreases for two reasons. First, some biological signal is contained in the discarded principal components. Second, models with a biological top feature still use technical artifacts and generate improved predictions as a result (**Fig. 10b**). Even if a model is not explicitly using a technical artifact, it may use a biological feature which is associated with a technical artifact.

**Fig. 10:**
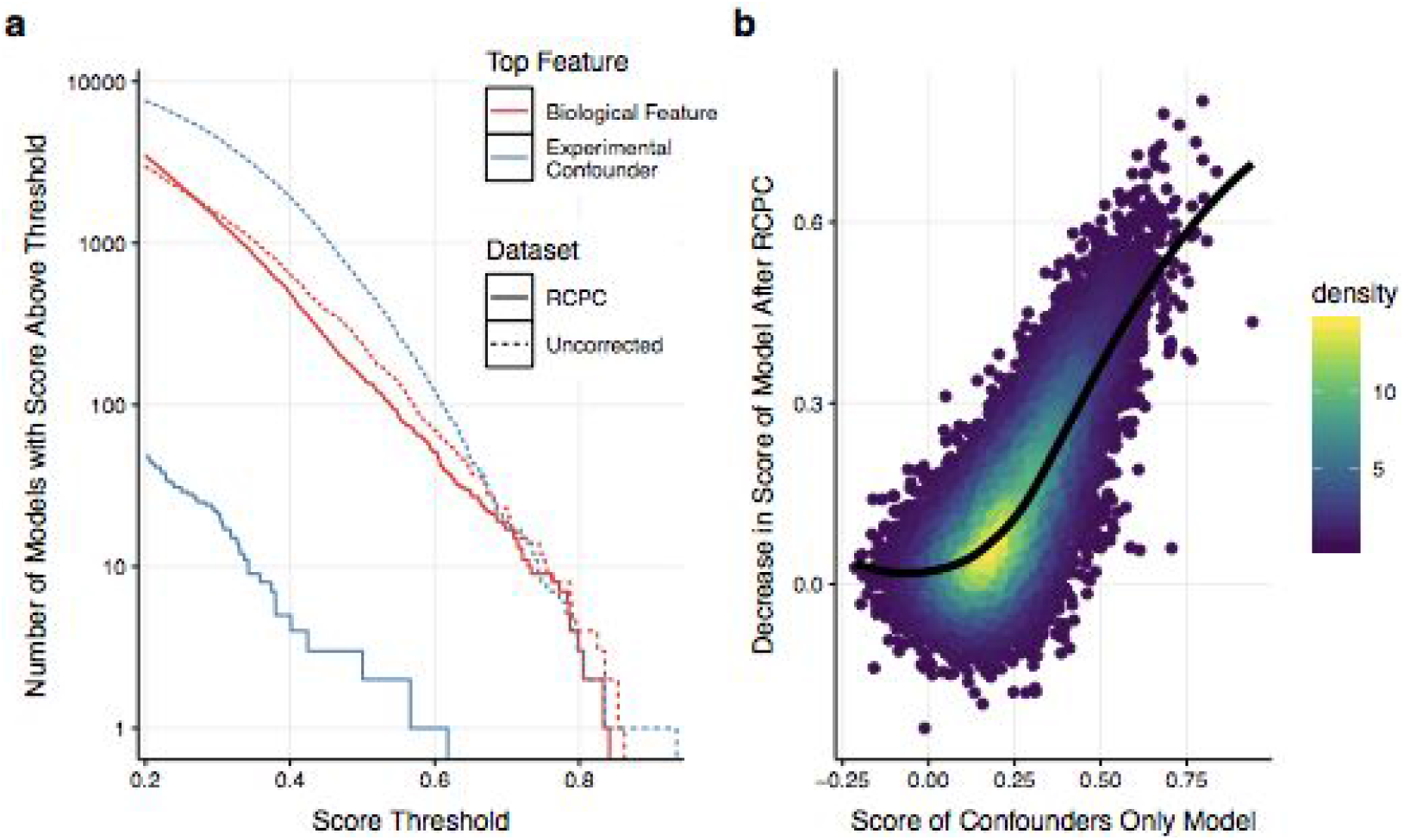
Effect on dependency prediction: (a) Survival curves for models trained to predict corrected and uncorrected gene dependency using CCLE cell line features and experimental confounders. Only models with score > .2 are shown. (b) Relationship between the decrease in the predictability of a gene after RCPC (using biological features and experimental confounders) and the predictability of a gene using only confounding features.

## Post-Processing

### Generating Dependency Probabilities

The majority of CERES scores appear to follow the distribution of known nonessential or unexpressed genes, suggesting that the true cell viability impact of knocking out these genes is zero (**Fig. 11a**). On the other hand, if knocking out a gene produces any nonzero reduction of viability, cells with that gene knocked out will vanish as a proportion of the cell pool over a sufficiently long time interval. Therefore it may sometimes be less interesting to ask how strongly depleted cells are after some gene knockout and more interesting to ask how likely it is that the observed depletion represents a real viability phenotype rather than noise.

**Fig. 11:**
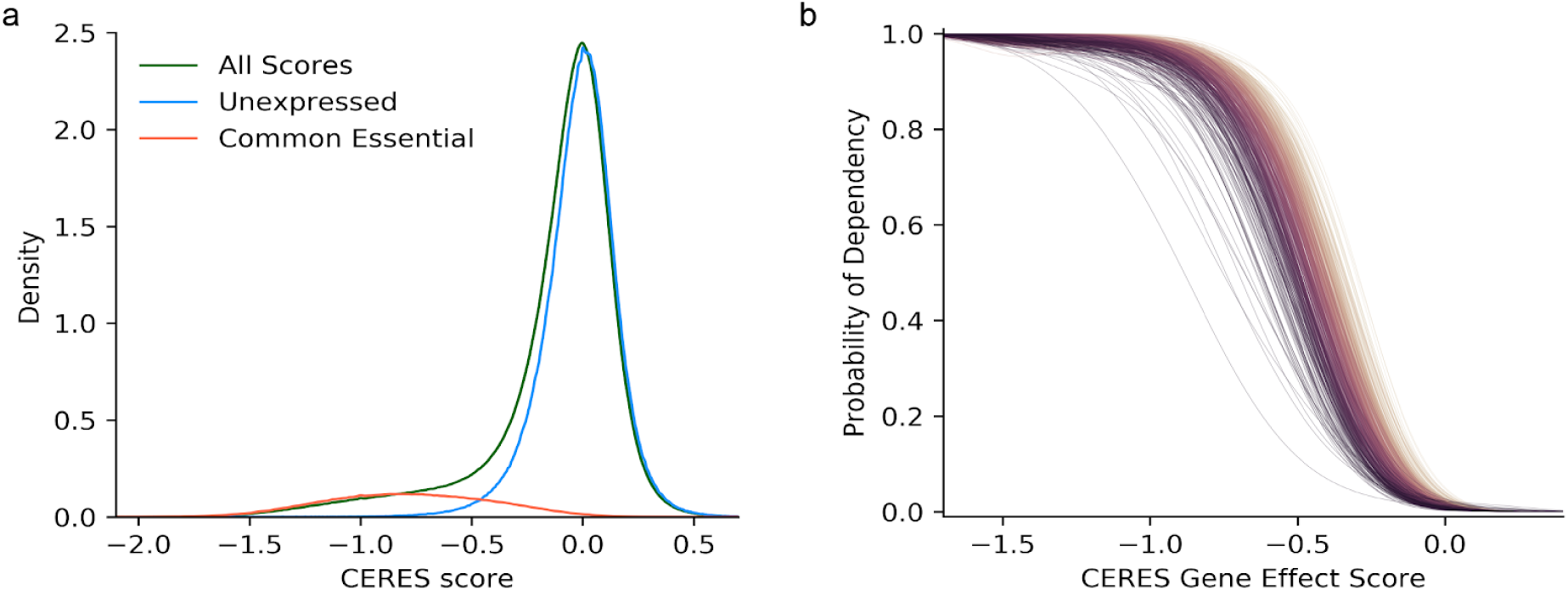
CERES scores and the probability of dependency: (a) Distribution of CERES gene scores for three different groups of score. The unexpressed gene distribution has been scaled by 0.84 and the common essential distribution by 0.12. (b) Relationship between the dependency probability and gene effect score. Each plotted line represents a different cell line. Lines are colored by the area under the curve.

To address this question, for each gene effect score in each cell line we estimate the probability that it represents a depleting phenotype. Specifically, we decompose the distribution of CERES scores in each cell line into the combination of two distributions: a null distribution representing no viability effect, and a positive distribution representing a real viability effect from a gene knockout. Both distributions are fit nonparametrically. The null is given by the distribution of the scores of unexpressed genes (those less than 0.02 in CCLE TPM-processed RNA-Seq expression data) smoothed with a Gaussian kernel of width 0.05. For cell lines with no available expression data, the Hart nonessentials are used instead. Beginning with the DepMap 19Q3 data release, positive distribution is similarly derived from the distribution of the positive control gene scores (given by *common_essentials.txt*, see Choice of Gene Controls) in the cell line. Previously, we used the essentials found from the Avana data itself and provided in *Achilles_common_essentials.txt*; the change in estimated probabilities using either set of positive controls is minimal. For computational efficiency, the probability density function (PDF) of each distribution is precomputed over a grid of points spaced at intervals of 0.01.

Both distributions are long-tailed, and it is necessary to avoid artifacts due to the low density of points at the extremes of each distribution. This is accomplished by adjusting the null PDF so that points right of zero with density less than 10^−16^ are assigned density of 10^−16^, and similarly for the positive PDF so that it is at least 10^−16^ at all points to the left of −0.8. Although this procedure would ordinarily cause its own difficulties, in our case we are only ever interested in the ratio of the probability densities, and this is well-behaved after forcing the tails to have a small residual density. The grid of values for each PDF is then resmoothed using a gaussian kernel of 0.15 width. The null and positive probability densities are then computed for each gene score for the cell line using linear interpolation.

We then employ a standard Bayesian E-M optimization procedure with a single free parameter: the total fraction of gene scores in the cell line which was generated by the null distribution. We use an initial guess of 0.85, which is generally close to the final result. The final probability of each gene score being generated by the positive (real depletion) distribution is provided to users in the matrix *Achilles_gene_dependency.csv*.

The relationship between dependency probability and CERES gene effect is monotonic and approximately sigmoidal for all cell lines, but the dependency probability associated with intermediate gene effect scores varies widely between lines (**Fig. 11b**). This is due to the variance in screen quality between different lines. In a low-quality screen, all dependency probabilities are flattened towards the prior (generally around 0.15), creating a shallower slope. Consequently, a CERES score of −0.5 has very different interpretations in high- and low-quality screens.

### Identifying common dependencies

The large variety of cancer cells that have been assayed in Achilles can be exploited to identify genes which are universally important for the viability of human cells. We provide a list of such genes in the file *Achilles_common_essentials.txt*, distinct from the prior known common essentials used elsewhere in the pipeline (which are provided in the file *common_essentials.txt*). To generate this list, our approach follows the basic intuition that if a gene is universally important for cell viability, it should fall in the top Z most depleting genes in at least 90% of cell lines. (We assume that noise or contamination might cause the gene not to score strongly in up to 10% of cell lines.) The threshold ranking quantile Z can be chosen in a data-driven way. For a given gene, we can rank its gene effect score in each cell line, then arrange cell lines in order of increasing gene effect score for that gene. For most genes, the 90th-percentile least dependent cell line will show little or no depletion, while common dependencies will still be depleted (**Fig. 12a**). This creates a bimodal distribution of gene ranks in their 90th-percentile least depleted lines, where the left peak contains common dependencies and the right peak the remainder of genes (**Fig. 12b**). Accordingly, we choose the threshold Z to be the point of minimum density between the two peaks using a Gaussian smoothing kernel with width 0.1 and rounding to the nearest 0.001. The resulting list of common essentials is insensitive to the choice of 90th-percentile cutoff (**Fig. 12c**) due to the change in the stringency of the threshold Z as the choice of percentile cutoff varies (**Fig. 12d**).

**Fig. 12:**
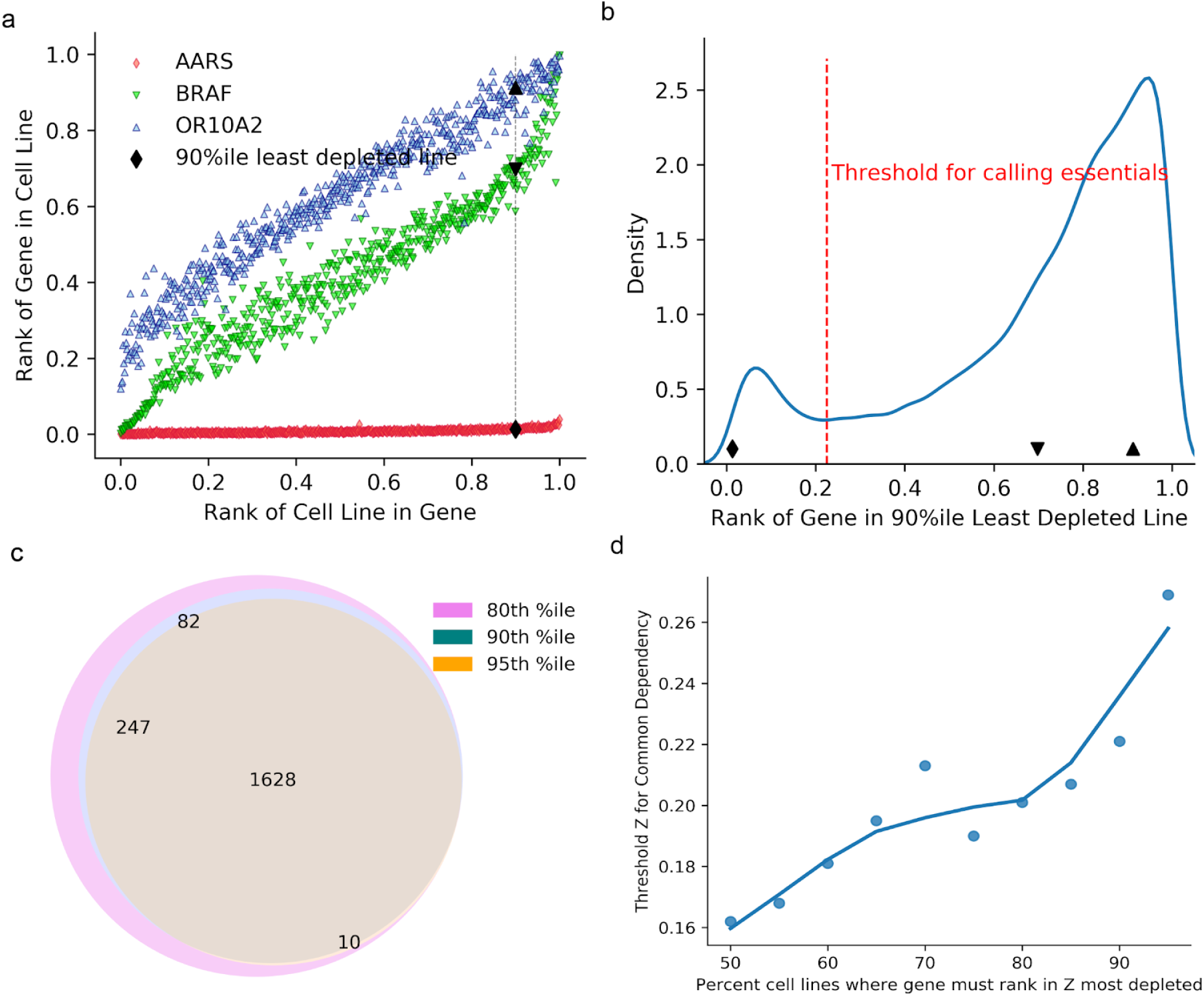
Identifying common essentials in CERES data: (a) For three example genes, the ranking of the gene’s CERES gene effect score among all genes in the cell line, with cell lines sorted in increasing order of CERES score for that gene. For each gene, its ranking in its 90th percentile least depleted line is highlighted. (b) Distribution of gene scores in their 90-percentile least depleted lines. (c) Overlap of common essentials when the same procedure is performed as in (b), but requiring 80% or 95% of cell lines to have the gene in the top X most depleting genes for those lines. (d) The threshold Z found at the minimum density of the distribution of gene scores in the given

## Analysis Methods

### Predicting Dependencies

We used scikit-learn’s RandomForestRegressor to predict gene effect scores and calculate feature importance from avana_public_19Q1 before and after the PC batch correction. The forest is constrained to have 100 estimators, maximum depth eight, and minimum samples per leaf node five. The following CCLE features were used:

- RNASeq expression data for both protein-coding and non-coding regions
- Mutation statuses, broken into three binary matrices:

∘ Damaging, true if the gene had a damaging mutation
∘ Hotspot, true if the gene had a nondamaging mutation in a TCGA or COSMIC hotspot
∘ Other, true if the gene had other conserving or nonconserving mutations
- Gene level copy number prioritized as described earlier in this manuscript
- Reduced representation bisulfite sequencing methylation of CpG islands and transcription start sites
- Histology as annotated in the Dependency Map portal

We also supplied the confounding factors of control separation, replicate correlation, pDNA batch, and growth media (**Supplementary Table 1**). Cell lines missing in any feature set were dropped, leaving 549 cell lines. Continuous features were z-scored. All remaining missing feature values were filled with zeros.

For each gene, training and prediction were performed over ten folds of cross-validation. For each fold, cell features were filtered to the 1,000 with highest Pearson correlation to the target gene effect profile in the training set, the forest was trained, and predictions were made and stored for the remaining 10% of validation cell lines. After all ten folds were completed, the complete set of out-of-sample predictions of gene effect were correlated with the actual gene effect profile to score the model. To report feature importance, the model was retrained on all samples.

### Replicate reproducibility – generating null distributions

Null distributions were generated by randomly selecting two replicates from all replicates and taking the Pearson correlation between them. This was done 1 million times for each null distribution.

### High variance genes

High variance genes were found by looking at the variance of gene effect across cell lines for each gene. The genes with the highest variance (top 3%) were taken; this yielded 529 genes.

### Predicting principal components with technical artifacts

PCA, with column variance normalization, indexed by sample/cell line, was performed on log-fold change data, gene effect data before cell line normalization, gene effect data after cell line normalization, and dependency probability data. Principal components were predicted by a linear model using the pDNA batch of each sample/cell line (one-hot encoded), the cas9 activity of each cell line, the nonessential z-score (recalculated for each level of data, calculated using the 19q3 updated positive control gene set)), and the replicate correlation of each sample/cell line (calculated at the log fold change level).

sgRNAs with NAs were discarded, and genes with variance equal to zero were removed before PCA. Cell lines with NA values for technical artifacts were dropped.

### RCPC data correction

PCA, with column variance normalization, indexed by sample/cell line, was performed on gene effect level data after cell line normalization and gene zscoring. The first six principal components and component loadings were removed and the truncated matrices were multiplied. The columns of the resulting matrix were multiplied by the original variance normalization factors and the columns means were added back. Finally, cell lines were renormalized according to the procedure in the CERES Normalization section.

### Linear regression-based correction

Each gene in gene effect level data, after cell line normalization, was predicted by a linear model using cas9 activity, nonessential z-score, pDNA batch and replicate correlation. Missing features were mean imputed. The residuals of the model were computed and the original gene means were added back to create the corrected gene effect matrix. Finally, cell lines were renormalized according to the procedure in the CERES Normalization section.

### ComBat correction

ComBat only supports the correction of data using discrete batches, but the cas9 activity and nonessential z-score of a cell line are continuous values. To create discrete batches, continuous quantities were split into 3 equal sized groups. ComBat was first applied to correct cas9 activity with nonessential z-score (discretized) and pDNA batch as covariates. ComBat was then applied to correct nonessential z-score with pDNA batch as a covariate. ComBat was then applied to correct pDNA batch without covariates. Finally, cell lines were renormalized according to the procedure in the CERES Normalization section.

The development of this procedure required multiple iterations. Two factors, the order in which experimental confounders were corrected and the size of continuous-variable groups, were varied extensively. ComBat procedures were compared based on their performance on the metrics summarized in Figure 7.

The SVA R package implementation of ComBat was used.

### RUV2 Correction

The random effect version of Naive RUV-2 implemented as naiveRandRUV in the RUVnormalize package was used. The ridge-regression regularization parameter was set to 0.1 and the rank of the estimated unwanted variation was set to 10. Nonessential genes were used as negative control genes.

Before application of the correction, gene means were removed. After application, gene means were added back and cell lines were renormalized according to the procedure outlined in the CERES normalization section of this report.

### Evaluation of cell line clustering

Each gene effect matrix was pruned so that every cell line had at least one other cell line of the same primary tissue type. Subsequently, *t*-distributed stochastic neighbor embedding (t-SNE) with perplexity equal to five was applied to each pruned gene effect matrix to reduce the dimension to two. K-means clustering was then applied to each embedded matrix with *k* equal to the number of primary tissue types represented. The adjusted Rand index was calculated between the k-means groupings and parent primary tissue groups. This process was repeated 100 times for each correction.

### Evaluation of correlation significance between related gene pairs

Correlation matrices for each gene effect matrix were computed and values within columns were ranked from high to low. The p-value for each correlation was equal to the rank of the correlation divided by 17633 (the number of other genes in gene effect level data). Q-values were then calculated across all p-values within a correction method. The number of biologically related gene-pairs with a q-value less than 0.1 were reported. Two lists of biologically related genes were used, human core complexes from CORUM and Human paralogs from Ensembl.

### Evaluation of concordance with gene expression data

The correlation between gene effect data and Public 18Q4 CCLE TPM expression data was evaluated according to the following procedure:

A pseudo count of 1 was added to Public 18Q4 CCLE TPM expression data and the resulting matrix was log2-transformed. All genes not present in the Public 18Q4 CCLE TPM and avana_public_19Q1 were discarded from both datasets.

Correlation matrices were computed between each gene effect matrix and the processed expression data (rows correspond to gene effect data and columns correspond to expression data). P-values were determined for each gene as the within row rank of the correlation between its knockout profile and its expression profile divided by 17395 (the number of genes in both datasets). Q-values were calculated for all genes within a correction method. The number of genes with a q-value less than .1 was reported.

### Evaluation of consensus biomarkers

Probabilities of dependency were calculated for DEMETER2^19^ gene scores using the methodology described under Post-Processing above. The dependency prediction pipeline described in the Analysis Methods section was run on DEMETER2 probabilities for genes with at least 10 dependent and 10 nondependent lines, in addition to Avana dependency probabilities. Genes with models from both datasets that shared a feature in common among the top two most important features and had model scores greater than 0.4 were selected. Spearman correlations were calculated between the corrected gene effect profiles of the selected genes and their top predictive features. Public 18Q4 CCLE TPM expression data and 18Q4 public DepMap Mutation call data were used.

### Implementation of ncPCA

A list of olfactory receptor genes was downloaded from the HGNC website (https://www.genenames.org/data/genegroup/#!/group/141). Genes with guides which targeted multiple genes as indicated in the guide_gene_map.csv file were dropped to create a list of 278 olfactory receptor genes.

Permutation testing was implemented as explained in Boyle *et al*.^17^. The first four principal components explained a significant percent of the variation among olfactory genes.

PCA was performed on the olfactory gene effect matrix (278 olfactory genes by 558 cell lines) using the *prcomp* function in R, without column scaling. Gene effect profiles were projected onto the first four principal components and returned to their original space via matrix multiplication, as specified in Boyle *et al*.^17^, to create a correction matrix. The difference between the original, full dataset and the correction matrix was used as the corrected dataset.

To generate a modified ncPCA dataset (benchmarked in Figure 7) we modified in the original protocol in two ways. First, gene means were removed before the correction procedure and added back after. Second, cell lines were renormalized after the gene means were restored.

More specifically, the normalization was performed in four steps. First, gene effect data was gene-centered and transposed (genes are rows and cell lines are columns). Second, permutation testing was performed to find principal components of nonessential genes that contain significant variation related to nonspecific killing. Third, the full, preprocessed gene effect matrix was projected onto the loadings of the nonsignificant principal components of the nonessential gene effect data and subsequently projected back to the original space. Lastly, gene means were then added to the matrix and cell lines were recentered according to the procedure in the CERES Normalization section.

### Correlation network significance testing

Significance testing of correlation networks for CORUM protein complexes was performed as described in Pan *et al*., 2018^18^, with the following modifications: 1) The results of 2000 (rather than 10,000) shuffled networks were used to generate the null distribution for significance testing; 2) Correlation ranks were computed on the full correlation matrices (17,634 × 17,634 genes) for each underlying dataset.

### Correlation network visualization

Four genesets defining essential protein complexes were taken from the GO Cellular Component Ontology as defined in the MSigDB C5 geneset collection (“GO_PROTEASOME_COMPLEX”, “GO_RNA_POLYMERASE_COMPLEX”, “GO_EXOSOME_RNASE_COMPLEX_”, “GO_CATALYTIC_STEP_2_SPLICEOSOME”). Genes present in more than one geneset were discarded. For each of the three gene effect matrices considered (uncorrected, RCPC and ncPCA-corrected), Pearson correlations were calculated between all pairs of genes present in the union of these four genesets. For each dataset, the top 400 correlations were taken as edges in a correlation network, which was visualized in the Cytoscape desktop application. The network layout was computed by combining the top edges from all three networks into a single combined network (1200 edges total), selecting the “Group Attributes Layout”, and selectively plotting edges specific to each dataset. Finally, to generate the ROC curve, true positives were defined as correlations between genes from the same geneset, while false positives were those between genes from different genesets.

## Acknowledgements

This work was funded by grants U01 CA176058 and U01 CA199253 and by the HL Snyder Foundation. The authors appreciate useful conversations with Julian Hess.

